# The prefrontal cortex outputs to the amygdala facilitate threat-discrimination learning

**DOI:** 10.64898/2026.06.26.734929

**Authors:** John H. Speigel, Tyler W. Bailey, Johannes Mayer, Edward Korzus

## Abstract

The medial prefrontal cortex (mPFC) plays a significant role in modulating the threat response, particularly in ambiguous circumstances. The mPFC performs this role through its connectivity with multiple brain regions, including the amygdala, long regarded as the central hub for threat responses. However, the roles of specific prefrontal projections to the amygdala in contextual threat discrimination are not yet fully understood, particularly regarding more complex learning tasks and when disentangling the functionally distinct prelimbic (PL) subunit of the mPFC. Here, we challenged mice with a contextual differential threat conditioning (DTC) learning task in which subjects were repeatedly exposed to one context predictive of a foot shock (CS+) and to a similar yet distinct context that was not (CS-). While control mice showed a similar threat response in both contexts immediately after threat conditioning, within a few days of contextual exposures, controls acquire threat discrimination and freeze less to CS- than to CS+ during late DTC. However, we found that inducing localized hypofunction of neuroplasticity in PL neurons projecting to the basolateral amygdala (BLA) impairs performance on DTC. This finding identifies the specific population of neurons in PL cortices as a critical site for learning to discriminate threat.

## Introduction

Threat discrimination refers to the cognitive or systemic ability to correctly distinguish between real threats and harmless stimuli. An impairment in this ability is often associated with anxiety disorders and post-traumatic stress disorder (PTSD). Inadequate initiation of threat-avoidance or defensive behaviors can result in injury or mortality. Conversely, the unwarranted enactment of these behaviors may lead to inefficient energy expenditure or missed opportunities to obtain essential resources.

Appropriate threat responses can be acquired in threat discrimination learning, which involves learning the correct prediction of cue–outcome relationships such as cue-threat, cue-safety, or cue-neutrality contingencies. To investigate the neural foundations of the modulation of threat-related processes, numerous prior studies have effectively utilized cued threat conditioning. This methodology involves classical conditioning in which subjects are trained to associate a neutral stimulus, such as an auditory tone, with an aversive unconditioned stimulus (US), thereby eliciting a threat response solely to the cue ^1, 2^. However, the evidence on which animals must make threat/safety judgments in naturalistic circumstances is typically much more subtle and complex than the cues used in cued threat conditioning and may require different neural circuitry for processing^3^. An alternative approach that better replicates these natural challenges is contextual threat conditioning, a variant of threat conditioning in which the subject’s overall environment predicts whether they will experience the US. Contextual threat conditioning studies have identified both similar and distinct elements of underlying neural circuitry compared to those involved in cued threat conditioning ^3, 4^. For example, while the amygdala is critical for the acquisition and expression of both cued and contextual threat conditioning ^5, 6^, the hippocampus appears to be involved in the acquisition and expression of contextual, but not cued, threat conditioning^5, 7–9^.

Optimal threat-response modulation behavior depends on the ability to learn a threat-association and later modify that threat in response to additional information. Modulation of threat response, such as threat extinction and threat discrimination learning, depends on different neural circuits and brain regions than those that enable the initial acquisition of conditioned threat ^10, 11^. Extinction learning is the learned suppression of a conditioned threat response. It operates by forming a new memory rather than through the actual erasure of the original threat-association memory ^12, 13^. Understanding the neural circuitry of threat discrimination learning is relevant to understanding the etiology of human anxiety disorders such as post-traumatic stress disorder (PTSD) and generalized anxiety disorder, both of which manifest an inability to suppress threat responses ^14–18^.

The amygdala is an essential brain region for most forms of threat-related learning and behavior ^19–26^. The amygdala integrates threat-related information from across the brain, serves as a major locus for threat-association memory storage, and triggers threat-response behaviors through its outputs to effector regions. In addition, studies have increasingly indicated that the medial prefrontal cortex (mPFC) is another region with major involvement in learning and modulating the threat response, largely through its reciprocal connectivity with the amygdala. In particular, bidirectional connectivity between mPFC and the basolateral amygdala (BLA) is prominent ^33, 34^. It has been implicated as one of the primary routes by which the mPFC affects the modulation of threat response learning ^35, 36^. The modulatory influence of the mPFC appears important for safety learning processes such as threat extinction and threat discrimination learning^37, 38^.

Although some studies have examined the entire mPFC as a single functional unit, other work has identified distinct roles for its prelimbic (PL) and infralimbic (IL) subdivisions in threat learning. PL has been implicated as the primary circuit underlying the mPFC’s role in strengthening threat-association^3, 28, 29, 31, 39^. Conversely, IL has been interpreted as primarily involved in the extinction of conditioned threat ^28, 40^ ^41^, and threat discrimination learning ^42^. However, significant gaps remain in our understanding of the full roles of PL in threat discrimination learning, particularly regarding the functional roles of subregion-specific projections (such as PL projections to BLA) in contextual threat response modulation ^29^. In the present study, we tested the putative role of PL neurons projecting to the BLA as a locus for contextual threat-discrimination learning by disrupting the neuroplasticity of this neuronal subpopulation in mice. We found that this manipulation impairs threat discrimination learning. This finding demonstrates that PL neurons projecting to the BLA constitute a learning site for modulating contextual threat responses.

## Methods

Subjects: The UC Riverside Institutional Animal Care and Use Committee approved all procedures in accordance with the NIH guidelines for the care and use of laboratory animals. 28 total male C57BL/6 mice (14 per group) from Taconic were used for this experiment. Subjects were approximately three months old (P90) during surgery. Mice had ad libitum access to food and water and were maintained on a 12/12 h light/dark cycle. All behavioral experiments were performed during the light phase of the cycle. Old bedding was exchanged for fresh, autoclaved Sani-Chips bedding each week. Before the start of the experiment, subjects were housed in three or four animals per cage with same-sex littermates. From the start of surgeries until sacrifice, subjects were housed with a single ovariectomized female companion mouse (which was not subjected to any experimental procedures) to avoid isolation stress. Tissues were harvested postmortem.

### Surgeries (virus injections)

In order to selectively manipulate the PL neurons projecting to BLA, all subjects received injections of anterograde AAV1.CaMKII.GFP-Cre into bilateral PL and of retrograde HSV.hEF1a.LS1L.CBPΔHAT.mCherry (for experimental subjects) or HSV.hEF1a.LS1L.mCherry (for controls) into bilateral BLA. Injection surgery procedures were based on those described by Cetin et al. (2006). All syringe movement (including virus aspiration and ejection) and skull calibration were controlled by a StereoDrive robotic stereotaxic frame (Neurostar). Mice were placed into an isoflurane chamber to induce anesthesia, mounted on a heating pad and into the stereotaxic apparatus, and supplied with a constant flow of isoflurane/oxygen mixture adjusted as needed to maintain a surgical plane of anesthesia (verified by the absence of toe-pinch response). Rimadyl (Caprofen, 5 mg/kg) and Buprenorphine (0.1 mg/kg) were injected at the start of surgery to reduce pain and inflammation, and ophthalmic ointment was applied to prevent eyes from drying out. Sterile saline (0.9%; 0.01 mL/g) was injected hourly to prevent dehydration. The scalp was shaved and sanitized with Betadine (Povidone-iodine) followed by 70% isopropyl alcohol. A midline incision was made on the scalp, and surgical hooks were placed to keep the skull exposed. The head was leveled by adjusting the subject’s positioning while checking the stereotaxically-measured coordinates of skull landmarks until bregma and lambda were at equivalent dorsoventral positions to one another, as were the points 2 mm left and right of bregma. Injection sites were calculated relative to bregma coordinates, incorporating a scaling factor based on the distance between bregma and lambda, compared with the standard 4.2 mm. A handheld dental drill was used to thin the skull overlying the syringe paths to the injection sites. A 27G needle tip was then used to remove the thinned bone and underlying dura gently. Sterile saline (0.9%) was applied as needed to rinse away any blood and prevent the exposed cortex from drying out. The stereotactic syringe (Hamilton glass 10µL w. 33G beveled needle) was aspirated to take up an aliquot of working virus solution just before insertion to the first injection site. The smooth flow of the virus was confirmed under a dissecting microscope immediately before insertion at each injection site to ensure the absence of any obstructions. The syringe was inserted and withdrawn at a rate of 0.1 mm/second, and virus was mechanically ejected at a rate of 0.02 µL/min.

After needle insertion to target region, there was a 2-minute pre-injection waiting period to allow tissue acclimation before the injection began; then there was a 10-minute post-injection waiting period after the injection was completed before the needle was withdrawn, to facilitate uniform viral dispersal and minimize backflow. After all injections were completed, the craniotomies were sealed with bone wax and the scalp was sutured, and a second injection of Rimadyl was administered. Antibiotic ointment was applied to the scalp, and the subject was placed in an oxygen chamber until it regained alertness, then returned to its home cage. Postoperatively, subjects received Rimadyl injections (5 mg/kg) daily for two days and oral ibuprofen (0.1 mg/mL in drinking water) for two weeks and were monitored for any signs of distress or postoperative complications.

All viral injections were performed bilaterally, and all viral injections for each subject were performed sequentially within a single surgery. All injections using a particular virus were performed sequentially from a single aliquot of working virus solution; between the PL and BLA injections, the syringe was sterilized and cleaned with sequential rinses of 10% bleach, DI H_2_O, and 70% ethanol to prevent cross-contamination with different viruses. Stereotaxic coordinates for virus injection are given relative to bregma and are based on the Paxinos mouse brain atlas (Paxinos & Franklin 2001). For PL injections, 0.2 µL of AAV virus per hemisphere was injected to 2.1 mm rostral, ±0.37 mm lateral, and 2.1 mm ventral of bregma, approached at a 15° lateral angle relative to the dorsoventral axis. For BLA injections, 0.2 µL of HSV virus per hemisphere was injected to 1.7mm caudal, ±3.4 mm lateral, and 4.7 mm ventral of bregma, approached parallel to the dorsoventral axis. Angled injection paths were utilized for PL to reduce damage and backflow to ventral portions of mPFC. Four surgeries (on two subjects from each experimental group) were completed per day, spanning 7 days for all 28 subjects. Subjects had 7 to 13 days to recover from surgery before contextual threat response conditioning training began.

### Viruses

Aliquots of stock virus were stored at -80° C, while working concentrations of virus were stored at 4° C. Stock aliquots were thawed and diluted as needed so that no working virus was kept at 4° C for longer than 2 days before use. We used custom-made HSV viruses (MGH Core) and commercially available AAV variants from UNC viral vector core.

Viruses and working titers used were as follows:

AAV1.CaMKII.GFP-Cre (UNC Core); 5 x 10^12^ GC/mL.

HSV.hEF1a.LS1L.CBPΔHAT.mCherry (R.Neve, MGH Core); 4 x 10^7^ GC/mL.

HSV.hEF1a.LS1L.mCherry (MGH Core); 4 x 10^7^ GC/mL.

### Behavior

Behavioral equipment: Threat conditioning was performed in 12.0” L x 10.2” D x 12.0” H plexiglass threat conditioning chambers (Lafayette Instrument Co.) equipped with an overhead IR-capable video camera, speaker, IR light, chamber light, shock grid, and ventilation fan, all controlled by FreezeFrame 4 software (Actimetrics). Each chamber was placed within an opaque sound-attenuating box, and four such chambers housed in the same testing room were used to run four subjects at once through all behavioral trials. Chambers were cleaned immediately before each trial with 70% ethanol and DI water and disinfected at the end of each day with Quatricide. Chambers could be set up in three configurations possessing distinct visual, auditory, and olfactory cues: Contexts A, B, and NS. Context A consisted of unadorned chamber walls, a lemon extract scent cue, an auditory 0.5 Hz frequency-modulated upsweep (2 to 9 kHz), and shock grid flooring configured as a single layer of parallel bars. Context B consisted of chamber walls covered with laminated inserts depicting vertical black-and-white stripes, a vanilla extract scent cue, a constant 2.8 kHz auditory tone, and shock-grid flooring configured as two staggered layers of parallel bars. NS consisted of an open-topped 9” L x 6” D x 6.5” H plastic box containing a layer of Sani-Chips bedding (changed out for each subject) and placed in the center of the threat conditioning chamber, which was set up with unadorned walls and no scent or auditory cue. Subjects could not leave the inner box or contact the shock grid during NS exposures. Contexts A and B were designed to be similar yet distinct, while NS was intended to be substantially different from them and to serve as a neutral context. Contexts A and B were counterbalanced with respect to US association; so, for half of the subjects, Context A exposure always predicted US (foot shock, 2 s, 0.75 mA) except during context habituation, while Context B never predicted US; and vice-versa for the other half of the subjects. Throughout this article, the US-associated context is referred to as CS+ and the US-unassociated context as CS-, although this counterbalanced design means that the conditioning chamber configuration designated by either term was Context A for half of the subjects and Context B for the other half.

Behavioral testing (Fig. 2): Before the first context exposure, all subjects were handled (2 minutes each) once per day for three days by the experimenter performing the upcoming course on contextual threat conditioning (TC) and contextual differential threat conditioning (DTC). Then, for the two days before TC (Habituation phase), subjects were habituated to the three experimental contexts (CS-, CS+, and NS) by presenting each context once for 10 minutes, with no US administered. For TC training (day 1), subjects were exposed to three US (foot shock, 2 s, 0.75 mA) within a single 420 s context CS+ exposure, beginning after a 180 s baseline, with a 90 s inter-trial interval (ITI) and a 60 s poststimulus period. Over the next two days, subjects were exposed to NS twice daily for 242 s. 72 hours after TC, subjects underwent 6 days of training on DTC and each day mice were exposed for 242 s each to NS, then to CS- and CS+ in an order that alternated by day. CS+ exposures were always paired with US during DTC, while exposures to CS-and NS were never reinforced.

Behavioral analysis: Performance was analyzed semi-automatically using a video-based system controlled by FreezeFrame 4 software (Actimetrics). Subjects were recorded in grayscale at 30 frames per second during all context exposure trials, and the FreezeFrame software calculated the difference between consecutive frames by comparing the change of each pixel, producing an objective quantification of subject movement. Freezing was defined as any period of at least 1 second during which this movement value remained below an experimenter-configured freezing threshold. All freezing reported in figures (expressed as “% freezing”) describes the percentage of time spent freezing during the first 180 s of a context exposure before the application of any US (for threat conditioning, the freezing percentage is also reported for the 90 s ISI and 60 s post-stimulus).

### Histology

To localize viral expression and quantify the acetylation effects of CBPΔHAT, IHC was performed as previously described (Vieira et al. 2014). Mice were sacrificed with Nembutal (200 mg/kg, intraperitoneal injection) and transcardially perfused with 20 ml of PBS, followed by 20 ml of 4% PFA. Brains were extracted, soaked in 4% PFA overnight at 4°C, and then soaked in 20% sucrose at 4°C until they sank. The brains were then flash-frozen in embedding media (Tissue-Tek, 4583) using dry ice and ethanol before being stored at -80°C. Free-floating 40 μm coronal sections were sliced using a cryostat (Leica, CM1860) and stored in cryoprotectant (50% PBS, 30% ethylene glycol, 20% glycerol) at -20°C. Free-floating immunohistochemistry (IHC) was performed by washing sections two times for 10 min in 1x PBS followed by a 1 h incubation in blocking buffer (4% normal goat serum in washing buffer), then washing three times for 10 min in washing buffer (1x PBS w/ 0.3% Triton X7 100). The sections were then incubated overnight at 4°C in antibody diluent (2% normal goat serum in washing buffer) with primary antibodies. After three washes with washing buffer, the sections were incubated with secondary antibodies for 3 hr at room temperature in antibody diluent. The sections were then washed once with washing buffer for 10 min and then incubated with DAPI (Invitrogen, D1306, 1:1000) for 15 mins. Sections were then washed twice with 1x PBS for 10 min. Before imaging, sections were mounted on glass slides (Superfrost Plus, 12-550-15) using a mounting medium (ProLong Antifade, P36965). Primary antibodies used for viral expression mapping were chicken anti-GFP polyclonal antibody (Thermo Fisher, A10262, 1:2000) and rat anti-mCherry monoclonal antibody (Life Technologies, M11217, 1:2000),

Secondary antibodies used for viral expression mapping were Alexa488 goat anti-chicken IgG (Thermo Fisher, A-11039, 1:1000) and donkey anti-Rat IgG (Life Technologies, A18749, 1:1000).

The primary antibodies used for quantifying histone acetylation were chicken anti-GFP (Molecular Probes, 1:1000), anti-acetyl-Histone H3 (Millipore, 1:2000), and anti-acetyl-Histone H4 (Millipore, 1:2000).

Secondary antibodies used for quantification of histone acetylation were Alexa647-goat anti-mouse IgG, Alexa488-goat anti-chicken IgG, and Alexa647-goat anti-rabbit IgG (all Molecular Probes, 1:1000).

### Imaging

Coronal sections were imaged at 20x (20x/0.95 NA *XLUMPlanFl* objective) magnification using a semi-automatic laser-scanning confocal Olympus FV1000 microscope controlled by Olympus FV10-ASW software (v. 2.01). The gain and offset of each channel were balanced manually using Fluoview saturation tools for maximal contrast. All settings were tested on multiple slices before data collection, and all brain slices were imaged using identical microscope settings for data collection once established. Each channel was acquired in “Sequential Mode, Frame”. All images were acquired using the “Integration Type: Line Kalman” and “Integration Count: 2” to increase the signal-to-noise ratio. The background fluorescence was measured and subtracted for each image. The PL and IL subregions of mPFC and the BLA subregion of the amygdala were localized by overlaying images from the Allen Mouse Brain Atlas using ImageJ. Region of interest (ROI) analysis was used for quantification.

### Statistics and Data Analysis

Data are expressed as the Means ± SEM. N indicates the number of animals. Statistical analysis was performed using GraphPad Prism and Excel (Microsoft, Inc.). Student’s *t*-test or ANOVA was used for statistical comparisons. Pearson’s correlation (*r*) was used as an *effect size*. Where applicable, *post hoc* analysis with correction for multiple comparisons allows for error rate control as indicated. A p < 0.05 is considered statistically significant. Asterisks indicate statistical significance: * = p < 0.05, ** = p < 0.01, *** = p < 0.001, n.s. = not significant.

## Results

In order to investigate the role of PL-to-BLA projecting neurons in contextual threat discrimination memory, we performed bilateral injections of AAV virus expressing Cre recombinase (AAV1.CaMKII.GFP-Cre) into the PL region of mPFC, and of retrograde Cre-dependent virus (HSV.hEF1a.LS1L.CBPΔHAT.mCherry for experimental subjects or HSV.hEF1a.LS1L.mCherry for controls) into BLA. This injection design induces pathway-specific expression of CBPΔHAT (a transgene that impairs neuroplasticity through reduced histone acetyltransferase ability) in PL neurons projecting to BLA (Fig. 1A). Because subregion-specific characterization of functionality was an essential objective of this study, injection volumes and coordinates were carefully selected and refined through pilot surgeries to ensure injection target specificity. Post-mortem histological evaluation of our experimental subjects confirmed that we successfully achieved selective transformation of the PL subregion of mPFC, with robust GFP expression observed in PL but minimal spillover seen in IL or other adjacent regions (Fig. 1B); meanwhile, expression of HSV vector was localized to BLA, with minimal expression in adjacent regions of amygdala (Fig. 1D). Figure 1C shows PL neurons co-expressing GFP and mCherry, indicating successful dual infection of PL neurons projecting to BLA projection neurons (Fig. 1C).

**Figure 1.**
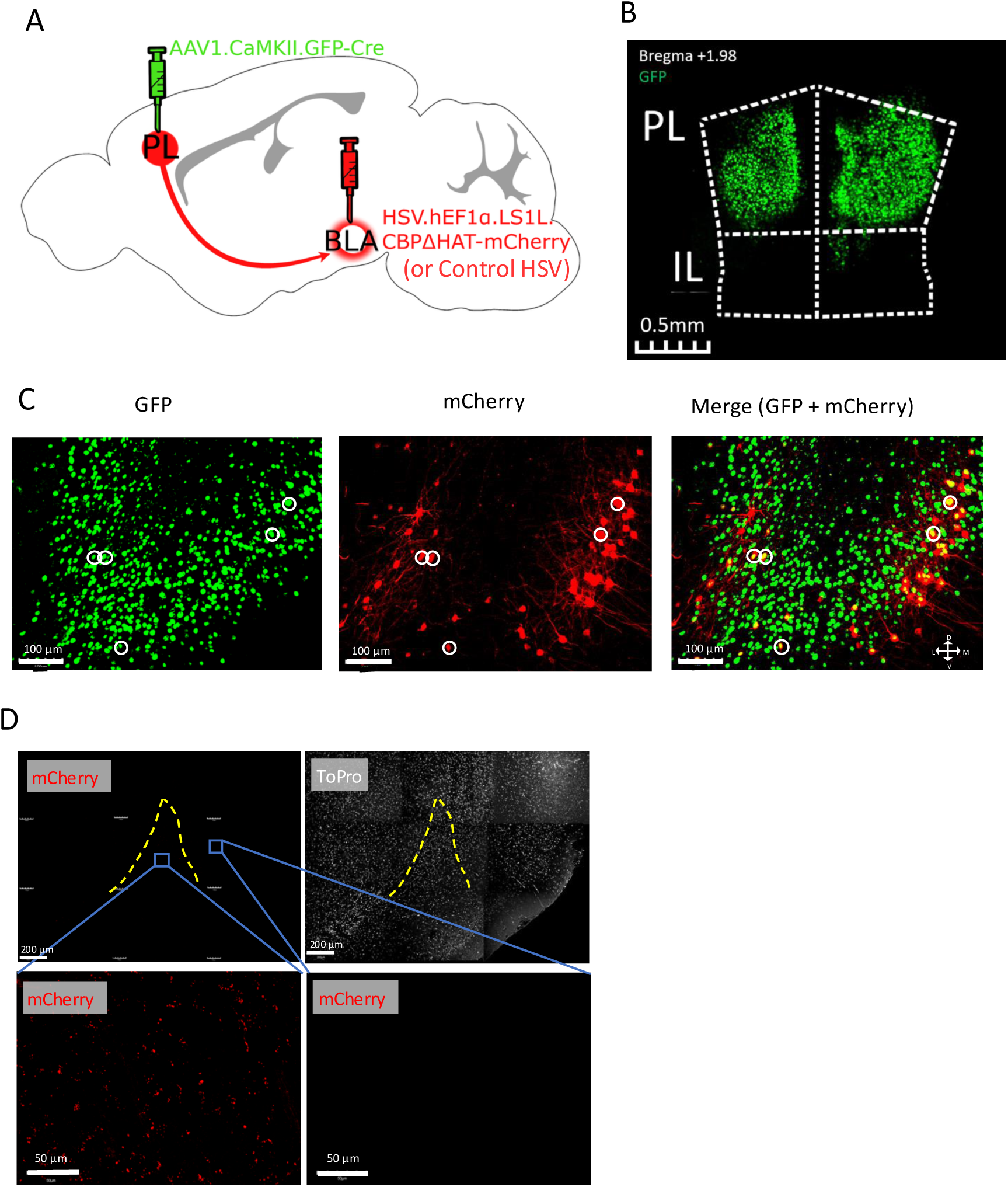
PL/BLA-CBPΔHAT mice selectively express CBPΔHAT in BLA-projecting PL neurons. **A:** Schematic illustration of viral injection strategy. PL injections of 0.3 µL of AAV1.CaMKII.GFP-Cre per hemisphere were delivered 2.1 mm anterior, 0.37 mm lateral, and 2.1 mm ventral to Bregma. BLA injections of 0.2 µL per hemisphere of HSV.hEF1α.LS1L.CBPΔHAT.mCherry (for experimental subjects) or HSV.hEF1α.LS1L.mCherry (for controls) were delivered 1.7 mm posterior, 3.4 mm lateral, and 4.7 mm ventral to Bregma. With this combination of vectors, only cells doubly infected by both anterograde AAV-Cre virus and retrograde HSV-CBPΔHAT virus express CBPΔHAT (and, consequently, experience a hypofunction of plasticity); and with this injection positioning, these CBPΔHAT-expressing cells are limited to the subset of PL neurons which project to targets in BLA (PL→BLA neurons). **B:** Representative mPFC (PL & IL) expression of AAV1.CaMKII.GFP-Cre. Transformed neurons are limited to PL, with minimal off-target labeling in IL. Coronal section, +1.98 mm from Bregma (landmarks based on Paxinos Mouse Brain Atlas). **C:** Representative PL expression of AAV1.CaMKII.GFP-Cre (GFP, green) and HSV.hEF1α.LS1L.CBPΔHAT.mCherry (mCherry, Red). The subset of PL neurons co-labeled with GFP and mCherry (merge) represents the targeted population of PL neurons that project to BLA. **D:** Upper-left: Representative expression of HSV.hEF1α.LS1L.CBPΔHAT.mCherry in PL-originating neuron terminals of BLA (composite image). Upper-right: Nonspecific ToPro nuclear staining of BLA. Lower-left: Expansion of central image of mCherry composite, depicting robust HSV transgene expression in BLA-projecting terminals. Lower-right: Expansion of center-right image of mCherry composite, depicting an absence of HSV transgene expression in regions outside of BLA. All histological samples were collected and analyzed after the completion of all behavioral testing.

We investigated effects on contextual threat learning and modulation by subjecting mice to a contextual threat conditioning (TC) and contextual Differential Threat Discrimination learning (DTC) protocols (Fig. 2). This task challenges subjects to learn to respond to threat context (CS+) followed by learning to discriminate between the CS+, similar but not the same non-aversive context (CS-) and substantially different neutral context (NS). After habituation to contexts (CS, CS-, and NS), contextual threat conditioning was performed through three US presentations in CS+. After assessing the freezing level during the first 180 seconds following mice’s placement in the training chamber (BL), the threat responses to CS+ were measured immediately after delivery of US1, US2, and US3 to assess short-term threat association recall during TC, and 72 hours after TC to assess long-term threat association memory recall (Figure 3A). We observed no differences in freezing behavior between CBPΔHAT and Ctrl and mice during contextual TC (Figure 3A; RM-ANOVA of Trial and Group: Trial x Group: F (2.679, 69.66) = 0.5, p = 0.66; Trial: F (2.679, 69.66) = 54.36, p<0.0001; Group: F(1, 26) = 0.57, p = 0.45). Sidak’s Post hoc multiple comparisons analysis indicated that both the Ctrl and PL-CBPΔHAT groups showed robust performance on the Threat Conditioning (Fig. 3A Dunnett’s multiple comparisons test, baseline (BL) vs. threat response after 3^rd^ US-CS+ Pairing: Ctrl, p < 0.0001; CBPΔHAT, p = 0.0001) and on the 72-hour threat-association recall (Fig. 3A. Dunnett’s multiple comparisons test, BL vs. 72-hours Recall: Ctrl, p < 0.0001; CBPΔHAT, p < 0.0001).

**Figure 2.**
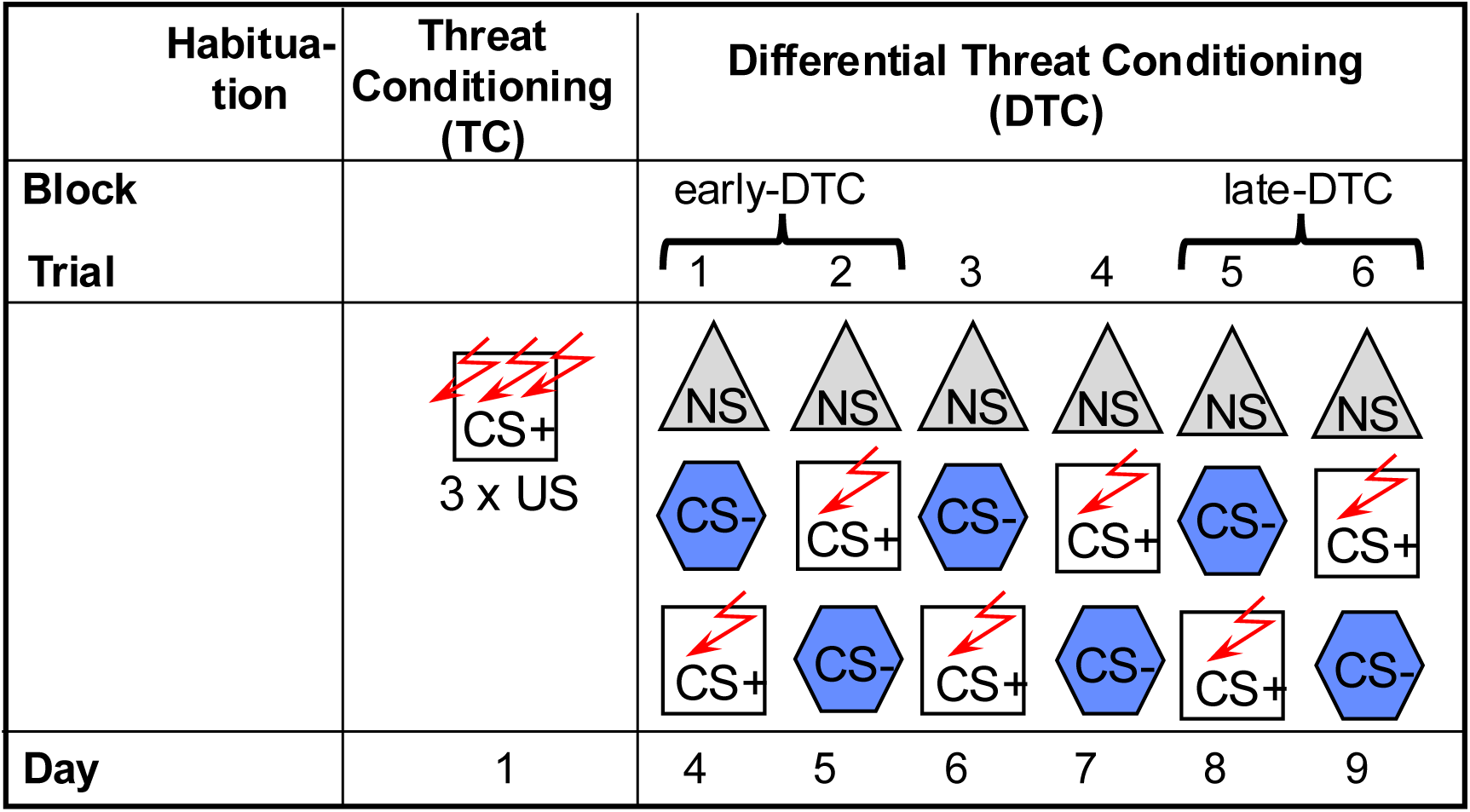
Contextual Threat Discrimination Learning (FDL) training schematic. Sequence of context exposures within each day is represented from top to bottom. CS+, aversive (US-paired) context; CS-, non-aversive (no-shock) context; NS = Neutral context (no-shock, dissimilar to CS- and CS+). The assignment of contextual cues (i.e., Context A vs Context B configuration) to CS- and CS+ was counterbalanced between subjects. Red arrows indicate CS-US pairings (2s 0.75 mA foot shock). As described in Methods, after 2 days of habituation to all contexts in the absence of US, subjects experienced 3 US-CS pairings during Threat Conditioning (Day 1; 180s prestimulus, 90s ITI, 60s poststimulus). 72 hours after TC, subjects were trained to behaviorally discriminate between CS- and CS+ through 6 days of Differential Threat Conditioning (DFC); each day consisting of an initial NS trial followed by CS- and CS+ trials presented in an alternating order (Days 4-9; 362s per trial; US delivered in CS+ after 180s prestimulus recording period for CS+ trials).

**Figure 3.**
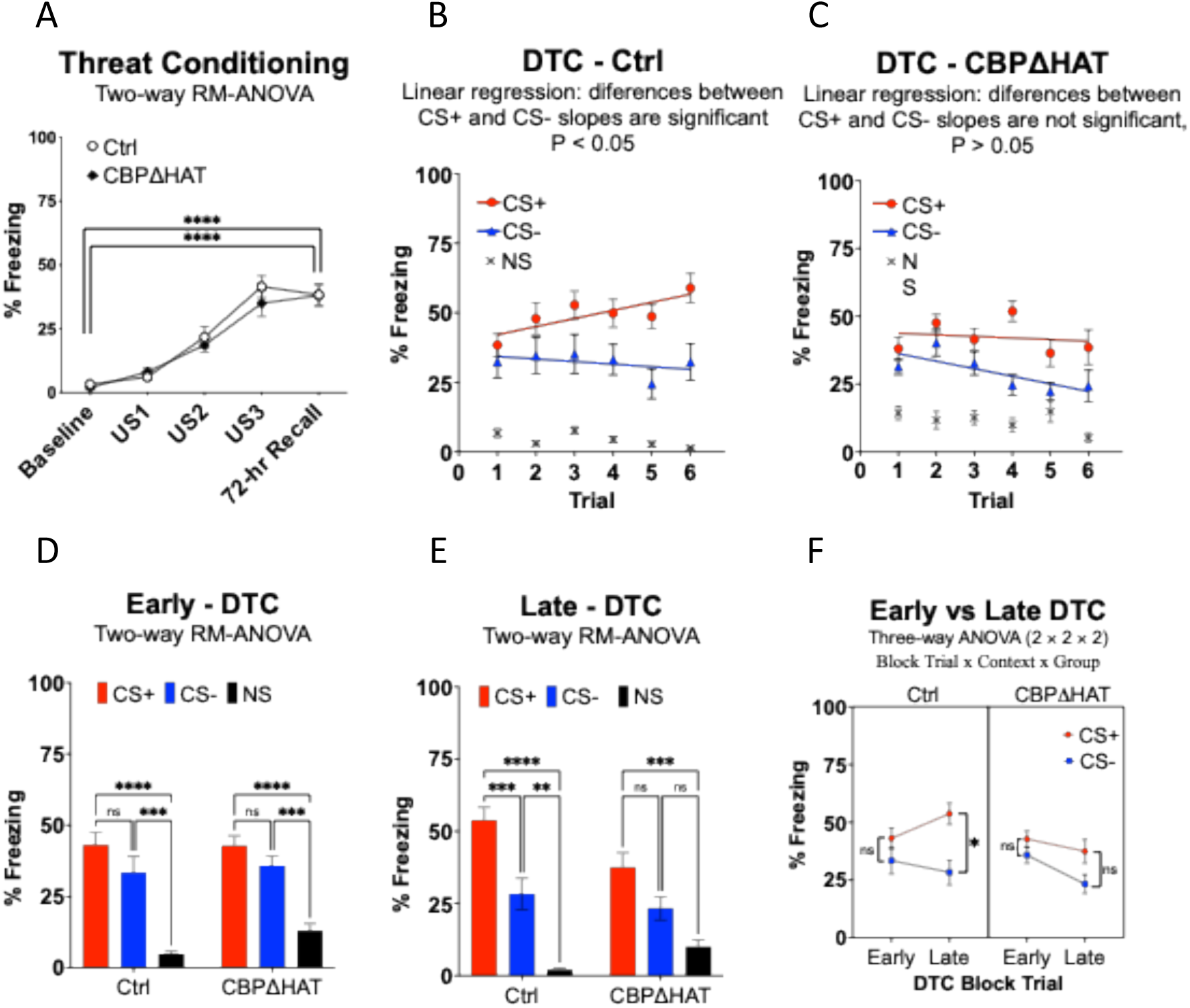
CBPΔHAT mice demonstrate normal acquisition of contextual threat conditioning but experience impaired contextual threat discrimination learning compared to controls. **A:** Both Control and CBPΔHAT mice robustly acquired conditioned threat to the US-associated context (CS+), with no significant differences between groups. **B:** Linear regression revealed that the CS+ response and the CS- response lines recorded on DTC in Ctrl are significantly different. **C**: Linear regression revealed that CBPΔHAT mice displayed a comparable CS+ response pattern to CS- responses during the entire DTC. **D:** During early-DTC, CBPΔHAT showed a similar CS+, CS-, and NS response pattern compared to Ctrl. **E**: During late-DTC, CBPΔHAT showed a deficit in DTC. **F:** Three-way ANOVA (matching by Block Trial and Context) of Block Trial x Group x Context confirmed impairment in DTC in CBPΔHAT compared to Ctrl *, p < 0.05, **, P < 0.01, ***, p < 0.001.

Three days after TC, subjects received six days of DFC to test their ability to learn to discriminate between CS+ and CS-(Fig. 2, 3B-F). A simple linear regression of responses to CS+ and separately to CS- was conducted to evaluate whether trial number predicted the difference in threat responses between CS+ and CS-. A comparison between the positive slope of the CS+ response line (y = 2.915 * x + 39.19) and the slope of the CS-response line (y = -0.93 * x + 37.88) in Ctrl mice demonstrated that the differences between slopes are statistically significant (Fig. 3B, F(1, 164) = 4.13, p = 0.04). In Ctrl mice, the positive slope of the CS+ response line is significantly greater than zero (F (1, 82) = 5.93, p = 0.0171), whereas the slope of the CS- does not significantly differ from zero (F (1, 82) =0.4, p = 0.53). Conversely, the comparison between the negative slopes of the CS+ response line (y = -0.59 * x + 44.35) and the CS- response line (y = -2.79 * x + 39.02) in CBPΔHAT mice revealed that the differences in the slopes are not statistically significant (Fig. 3B, F(1, 164) = 2.01, p = 0.16), thereby suggesting that projections from the prelimbic cortex to the basolateral amygdala are vital for threat discrimination learning. Notably, the slope of the CS+ response line in CBPΔHAT mice does not differ from zero (F (1, 82) = 0.2648, p = 0.61), while the slope of the CS-response in these mice is strongly descending (F (1, 82) = 7.043, p < 0.01). These data show that PL projections to the BLA are critical for the differential modulation of appropriate responses to both CS+ and CS- by enabling the enhancement of CS+ responses through experience with reminders of aversive experiences. Conversely, PL projections to the BLA are probably also responsible for the appropriate threat generalization of CS- responses by enabling cautious responses in ambiguous situations, as hypofunction targeted at these PL projections leads to faster safety learning.

To better understand how PL projections to BLA regulate environmental experience-triggered threat response modulation, we directly compared responses to different contexts during early and late DTC (Fig 2. Fig 3D-F) in CBPΔHAT mice and compared with Ctrl mice. Both Ctrl and CBPΔHAT mice performed the same during early-DTC, showing an inability to discriminate between CS+ and CS- during the initial two days of DTC (Fig 3D. two-way RM-ANOVA Group x Context: F (1.77, 46.08) = 1.002, p = 0.3665; Group: F (1, 26) = 0.7188, p=0.40; Context: F (1.772, 46.08) = 67.03, p < 0.0001. Ctrl: CS+ vs CS-, p > 0.05; CS+ vs NS, p < 0.05; CS- vs NS, p < 0.05. CBPΔHAT: CS+ vs CS-, p = 0.05; CS+ vs NS, p < 0.05; CS- vs NS, p < 0.05). Ctrl mice have learned to discriminate between aversive and non-aversive stimuli during DTC and have shown the ability to discriminate among CS+, CS-, and NS at late-DTC (Fig 3E. two-way RM-ANOVA Group x Context: F (1.956, 50.85) = 5.52, p=0.007; Group: F (1, 26) = 1.26, p=0.27, Context: F (1.956, 50.85) = 58.81, p<0.0001. Ctrl: CS+ vs CS-, p < 0.05; CS+ vs NS, p < 0.05; CS- vs NS, p < 0.05. However, CBPΔHAT mice showed a deficit in learning on the DTC and could not discriminate between CS+ and CS- at late DTC (Fig. 3F. CBPΔHAT: CS+ vs CS-, p > 0.05; CS+ vs NS, p < 0.05; CS- vs NS, p > 0.05). On the contrary to Ctrl, CBPΔHAT mice are unable to distinguish between CS-and NS during late-DTC (Fig. 3F. CBPΔHAT: CS- vs NS, p > 0.05), confirming again (see Fig 3C) that Pl-to-BLA projections may control appropriate threat generalization. Three-way ANOVA (matching by Block Trial and Context) of Block Trial x Group x Context confirmed impairment in DTC in CBPΔHAT compared to Ctrl (Fig. F. Context: F (1.000, 26.00) = 25.56, p<0.0001; Block Trial: F (1.000, 26.00) = 2.47, p=0.1280; Group: F (1, 26) = 0.79, p=0.38; Block Trial x Group: F (1.000, 26.00) = 8.98, p=0.006; Block Trial x Context: F (1.000, 26.00) = 14.60, p = 0.0007; Block Trial x Group x Context: F (1.000, 26.00) = 1.570, p = 0.2214). Tukey’s multiple comparison test showed that both CBPΔHAT and Ctrl mice did not discriminate between CS+ and CS- during early DTC (CS+ vs CS- at early DFC: CBPΔHAT, p > 0.05; Ctrl, p > 0.05). Ctrl mice showed robust performance on the DTC task and displayed strong discrimination between CS+ and CS- during late DTC (p = 0.001), while threat responses to CS+ and CS- in CBPΔHAT were the same at late DTC (p = 0.32). In summary, while CBPΔHAT mice showed normal performance on TC (Fig. 3A), these mice exhibited impaired performance in contextual threat discrimination learning compared with Ctrl mice (Fig. 3B-F).

To confirm that CBPΔHAT expression was producing the expected reduction in histone acetylation in transformed neurons, after behavioral testing was completed, a random subset of subjects’ brains was stained for H3 or H4 histone acetylation, and the extent of acetylation was quantified. We indeed observed substantial reductions in both H3 and H4 histone acetylation. Acetylation levels were normalized relative to Ctrl subjects. Average histone H3 acetylation levels were substantially lower in CBPΔHAT mice than in Ctrl mice (Fig. 4, CBPΔHAT mice (Mean = 0.65 ± 0.11) vs Ctrl mice (Mean = 1.00 ± 0.08); unpaired t test, two-tailed, t(14) = 2.420, p = 0.03, effect size R^2^ = 0.30). In addition, average histone H4 acetylation levels were also substantially lower in CBPΔHAT mice compared to Ctrl mice (Fig 4B. CBPΔHAT mice (Mean = 0.67 ± 0.09) vs Ctrl mice (Mean = 1.00 ± 0.06); unpaired t test, two-tailed, t(14) = 3.2, p = 0.007, effect size R^2^ = 0.42).

**Figure 4.**
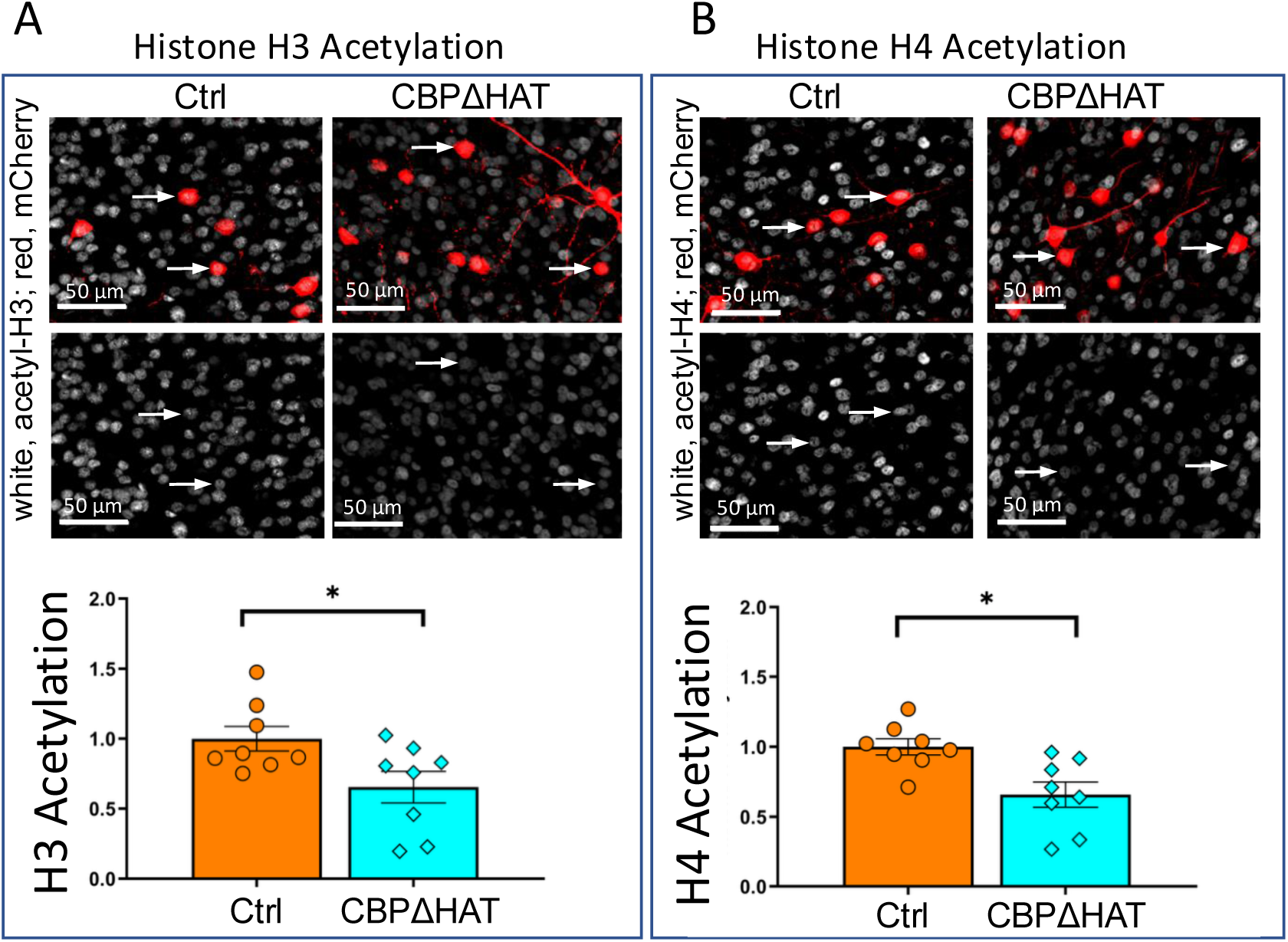
CBPΔHAT transformation induces reductions in both H3 (left box) and H4 (right box) histone acetylation. Top: Representative micrographs showing histological staining for both acetylation (white) and mCherry (red) in PL. Bottom: quantification of histone acetylation in Ctrl vs PL/BLA-CBPΔHAT mice, normalized vs Ctrl levels. Average histone H3 acetylation was substantially lower in CBPΔHAT mice vs Ctrl mice. Average histone H4 acetylation was also substantially lower in CBPΔHAT mice vs Ctrl mice. Ctrl, n=14; CBPΔHAT, n=14. * = p < 0.05; ** = p < 0.01. All histological samples were collected and analyzed after the completion of all behavioral testing.

This data indicates that BLA-projecting PL neurons serve as a significant locus for learning and control of threat-response modulation during DTC, as disrupting the neuroplasticity of this specific subpopulation is sufficient to impair the acquisition of threat discrimination between two similar contexts.

## Discussion

Using a contextual variant of a Pavlovian conditioning DTC task, we demonstrated the functional significance of PL neurons projecting to the BLA in contextual threat-discrimination learning. These data suggest that the neuroplasticity of PL neurons projecting to BLA is a crucial mechanism of that learning process. Our findings are consistent with prior evidence that PL is not essential for the initial acquisition or execution of simple conditioned threat response behavior but is heavily involved in more complex threat associations, such as those involving conflicting danger/safety environmental cues. Mice with the impaired plasticity of PL neurons projecting to BLA generated by targeted expression of CBPΔHAT mutant displayed typical acquisition and generalization of contextual threat, but showed an impaired ability to learn divergent behavioral responses to two similar contexts when only one is associated with an aversive outcome.

Before this study, the importance of the PL neurons projecting to BLA was implied by their anatomical prominence in PL-Amygdala connectivity. In initial studies, Vertes found that, in rats, PL’s prominent amygdala projections are concentrated in the BLA and the central nucleus (CeA) ^33^. It was also noted that PL to amygdala connections are stronger contralaterally than ipsilaterally and that PL’s projections to BLA and CeA originate more from the rostral than from the caudal region of PL. Cho et al. (2013) also found strong projections from PL to BLA in mice, particularly to the anterior portion of BLA ^36^. Kim et al. (2013) found that lesioning PL of rats that had learned a context-dependent threat association to a cue tone abolished their ability to exhibit greater threat response of the tone in the context in which it predicted a shock ^43^. However, such lesions did not affect the ability of rats that had learned a simple foreground contextual threat discrimination task to exhibit differential threat responses to the two contexts^43^. This latter finding suggests that the PL may not be required to retrieve contextual threat discrimination once learned. Current data indicate that long-term changes in PL do occur during the contextual threat discrimination process. Presumably, these changes enable PL to help entrain downstream regions, after which PL’s ongoing input is no longer essential for retrieval. At the population level, contextual threat conditioning has activated substantial engrams of PL neurons for both conditioned (CS+) and similar non-conditioned (CS-) contexts ^44^. There is also evidence that these engrams are used to accurately maintain and/or produce threat responses to CS+ contexts, as the extent of PL engram reactivation was positively correlated with the degree of contextual discrimination for CS+ engrams but not for CS- engrams ^44^.

## Conclusion

The present body of work contributes to this effort by identifying specific sites of contextual discrimination threat-association memory storage within the medial prefrontal cortex. Current studies have used a dual-virus injection CBPΔHAT transfection strategy to induce hypofunction of PL neurons projecting to the BLA and have observed impairment in contextual threat discrimination learning. Control mice could behaviorally discriminate between CS- and CS+ contexts within 6 days of training, whereas experimental mutant mice with epigenetic hypofunction targeted to PL neurons projecting to the BLA did not. Investigations of the role of the IL-to-BLA pathway and the function of specific projections from the BLA to subdivisions of the mPFC would be both intriguing and necessary for future research to understand the complexity of mPFC and amygdala interactions during learning to modulate threat responses in the presence of conflicting environmental cues.

## Funding

National Institutes of Health grant R01 MH106617 (EK)

Brain and Behavior Research Foundation grant (EK)

DoD/ARL grant W911NF-23-1-0145 (EK)

We thank undergraduate students Christopher Li, Ryan Wales, Alex Tran, Lindsay Melendez, Marie Sanchez, and Sandra Loza, whose efforts were instrumental to the success of this project.

